# RNA Polymerase II Independent Recruitment of SPT6 at Transcription Start Sites in *Arabidopsis*

**DOI:** 10.1101/506063

**Authors:** Chen Chen, Jie Shu, Chenlong Li, Raj K. Thapa, Vi Nguyen, Kangfu Yu, Zechun Yuan, Susanne E. Kohalmi, Jun Liu, Frédéric Marsolais, Shangzhi Huang, Yuhai Cui

**Affiliations:** Agriculture and Agri-Food Canada, London Research and Development Centre, London, Ontario N5V 4T3, Canada; Department of Biology, Western University, London, Ontario N6A 5B7, Canada; School of Life Sciences, Guangdong Provincial Key Laboratory of Plant Resource, Sun Yat-sen University, Guangzhou 510275, Guangdong, China; Agriculture and Agri-Food Canada, Harrow Research and Development Centre, Harrow, Ontario N0R 1G0, Canada; Guangdong Academy of Agricultural Sciences, Guangzhou 510640, China; Lead Contact

## Abstract

SPT6 is a conserved transcription regulator that is generally viewed as an elongation factor. However, emerging evidence show its potential role in the control of transcription initiation at genic and intragenic promoters. Here we first present the genome-wide occupancy of Arabidopsis SPT6-like (SPT6L) and demonstrate its conserved role in facilitating RNA Polymerase II (RNAPII) occupancy across transcribed genes. Further, we show that SPT6L enrichment is shifted, unexpectedly, from gene body to the transcription starting site (TSS) when its association with RNAPII is disrupted. Finally, we demonstrate that recruitment of SPT6L starts at TSS, and then spreads to the gene body during transcription. These findings refine the mechanisms underlying SPT6L recruitment in transcription and shed light on the role of SPT6L in transcription initiation.

## Introduction

It is well known that SPT6 is a transcription elongation factor, as evidenced by its physical association with elongating RNAPII (Andrulis et al., 2000; Kaplan et al., 2000; Mayer et al., 2010) and its ability to enhance elongation in vitro (Endoh et al., 2004) and in vivo (Ardehali et al., 2009). The Src homology 2 (SH2) domain of SPT6 recognizes and binds to phosphorylated serine 2 and tyrosine 1 repeats within the C-terminal domain (CTD) of RNA polymerase II (RNAPII), and to phosphorylated linker region preceding the CTD (Ardehali et al., 2009; Mayer et al., 2010; Sdano et al., 2017; Sun et al., 2010). Deletion or mutation of SH2 disrupts the interaction of SPT6 and RNAPII (Dronamraju et al., 2018; Mayer et al., 2010; Yoh et al., 2007) and significantly reduces the levels of SPT6 and RNAPII at transcribed regions of genes (Dronamraju et al., 2018; Mayer et al., 2010). Genetic and genomic studies in yeast have indicated the role of SPT6 and other elongation factors in control of intragenic initiation (Cheung et al., 2008; Hennig and Fischer, 2013; Kaplan et al., 2003). Recently, it was found that SPT6 was involved in regulation of genic initiation and mutation of SPT6 caused the reduced occupancy of TFIIB at genic promoters (Doris et al., 2018).

In *Arabidopsis*, there are two versions of SPT6: SPT6 (AT1g63210) and SPT6-like (SPT6L) (AT1g65440) (Gu et al., 2012). The transcript of *SPT6* was barely detectable in most of the tissues (Antosz et al., 2017) and no visible phenotype was observed in *spt6* mutants (Gu et al., 2012), suggesting that SPT6 may not play an essential role in transcription. *SPT6L*, however, appears to be commonly expressed (Antosz et al., 2017) and mutations in it led to the formation of aberrant apical-basal axis and embryonic lethality (Gu et al., 2012). Furthermore, SPT6L can be co-purified with RNAPII and other elongation factors (Antosz et al., 2017). These findings indicate its potential role in the regulation of transcription.

In this study, we examined the genome-wide occupancy profile of SPT6L and demonstrate its functional conservation in transcription elongation. By analyzing the global association between SPT6L and RNAPII, intriguingly, we found that the enrichment of SPT6L was shifted from the transcribed regions to transcription start sites (TSS) in the absence of its association with RNAPII. We further generated a series of domain deletions and showed that the HtH and YqgF domains of SPT6L are required for its TSS enrichment and even the distribution along gene bodies. Finally, we show that SPT6L was initially recruited at TSS and then spread to the gene body during transcription. In sum, our findings reveal novel mechanisms underlying the recruitment of SPT6L into the transcription machinery.

## Results

### SPT6L Co-occupies Genome-Wide with RNAPII over Highly Transcribed Genes

To gain insights into the functions of SPT6L in plants, we tagged the green fluorescent protein (GFP) to SPT6L (SPT6L-GFP) and introduced it into a *spt6l*^+/-^ (*SALK_016621*) heterozygous background. The transgene can fully complement the defects of the *spt6l* mutant (referring to *spt6l*^-/-^ homozygous mutant background hereafter) (Figure 1A to 1C) and the GFP signals were mainly detected in the nuclei (Figure S1A). Next, we profiled the genome-wide occupancy of SPT6L by chromatin Immunoprecipitation-sequencing (ChIP-seq) and found that SPT6L was mainly recruited to the transcribed regions of genes (Figure 1D). As SPT6 plays a key role in transcription elongation in other species (Ardehali et al., 2009; Endoh et al., 2004; Kaplan et al., 2000; Sun et al., 2010; Yoh et al., 2007), we examined the association of SPT6L with transcription in plants. First, we compared our SPT6L ChIP-seq data with published profiles of histone marks (Chen et al., 2017; Luo et al., 2013) and found that SPT6L binding genes were all marked with active histone modifications, but not the repressive ones (Figure S1B). Second, we profiled RNAPII occupancy in wild-type (WT) and *spt6l* mutants by ChIP-seq and compared with the SPT6L data. The ChIP-seq reads of SPT6L and RNAPII were highly correlated genome-wide (Figure 1E) and the occupancy of RNAPII was dramatically decreased in *spt6l* (Figure 1F). Third, the binding intensity of SPT6L is positively correlated with transcript levels (Figure 1G). These data indicate that Arabidopsis SPT6L likely plays similar roles in transcription as its homologs in other species.

**Figure 1:**
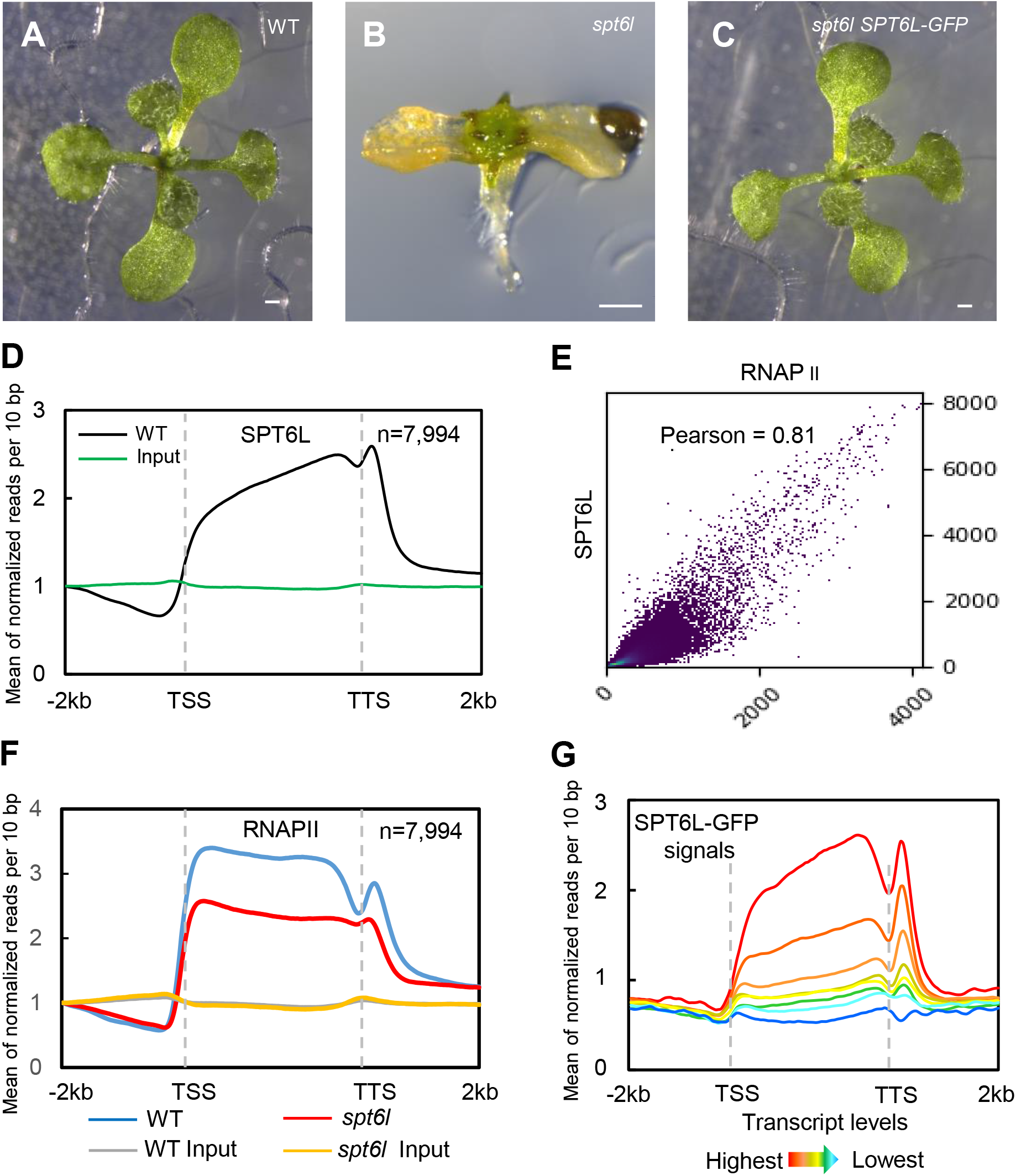
SPT6L is associated with transcribed genes and promotes transcription. (A) to (C) 14 days old WT (A), *spt6l* (B), and *spt6l* ProSPT6L:SPT6L-GFP (C) seedlings. Bar = 0.5 mm (D) Mean density of SPT6L occupancy at the SPT6L bound genes. Plotting regions were scaled to the same length as follows: 5’ ends −2.0 kb to transcription starting site [TSS]) and 3’ ends (transcription termination site [TTS] to downstream 2.0 kb) were not scaled, and the gene bodies were scaled to 3 kb. The y-axis represents the means of normalized reads (1x sequencing depth normalization) per 10 bp non-overlapping bin, averaged over two replicates (SPT6L) or one replicate (Input). Gene number (n) is indicated. (E) Reads correlation between SPT6L and RNAPII ChIP-seq (Pearson correlation value was indicated). The entire genome was equally divided into 100 bp non-overlapping bins and the numbers of reads were averaged over two replicates. (F) Mean density of RNAPII occupancy in WT and *spt6l* at SPT6L bound genes. Plotting regions were the same as in Figure 1D and y-axis values were the means of normalized reads (1x sequencing depth normalization) per 10 bp non-overlapping bin, averaged over two replicates or one replicate for RNAPII in *spt6l.* (G) Mean density of SPT6L occupancy in gene groups with different transcription levels. All *Arabidopsis* genes were clustered into 8 groups based on their transcript levels, from high to low, The y-axis values were the means of normalized reads (1x sequencing depth normalization) per 10 bp non-overlapped bin, averaged over two replicates.

### SPT6L Enriched at transcription start sites in the Absence of Its Association with RNAPII

In yeast, SPT6 interacts with both the phosphorylated C-terminal domain (CTD) of RNAPII and the phosphorylated linker region preceding the CTD via its SH2 domain (Ardehali et al., 2009; Sdano et al., 2017; Sun et al., 2010). Disassociation of SPT6 and RNAPII caused by the deletion of the SH2 domain significantly reduced the level of SPT6 and RNAPII occupancy along genes (Dronamraju et al., 2018; Mayer et al., 2010), but low levels of truncated SPT6 can still be detected at the transcribed regions of genes (Dronamraju et al., 2018; Mayer et al., 2010), suggesting the existence of RNAPII independent mechanism for SPT6 recruitment. To examine whether plant SPT6L also can be recruited to genes in an SH2 independent manner, we made an SH2 deleted version of SPT6L tagged with GFP (SPT6LΔSH2-GFP) and introduced it into *spt6l*^+/-^ (Figure S2A). Due to the lack of antibodies that can recognize the phosphorylated linker region of RNAPII, we chose to use the phosphorylated serine 2 of CTD (RNAPIIS2P) as an indicator of the active form of RNAPII. We then performed co-immunoprecipitation (Co-IP) experiments with the transgenic line and found that the deletion of SH2 indeed impaired the interaction between SPT6L and RNAPIIS2P (Figure 2A). Intriguingly, unlike the severely defected *spt6l* mutants (Figure 2B to 2D), the *spt6l* SPT6LΔSH2-GFP seedlings can grow bigger and develop small true leaves (Figure 2E to 2G). In line with the morphological phenotype, the introduction of SPT6LΔSH2 also partially rescued the genome-wide occupancy of RNAPII (Figure 2H and 2J). These findings indicate that SPT6LΔSH2 still retains some capacity in facilitating RNAPII transcription.

**Figure 2:**
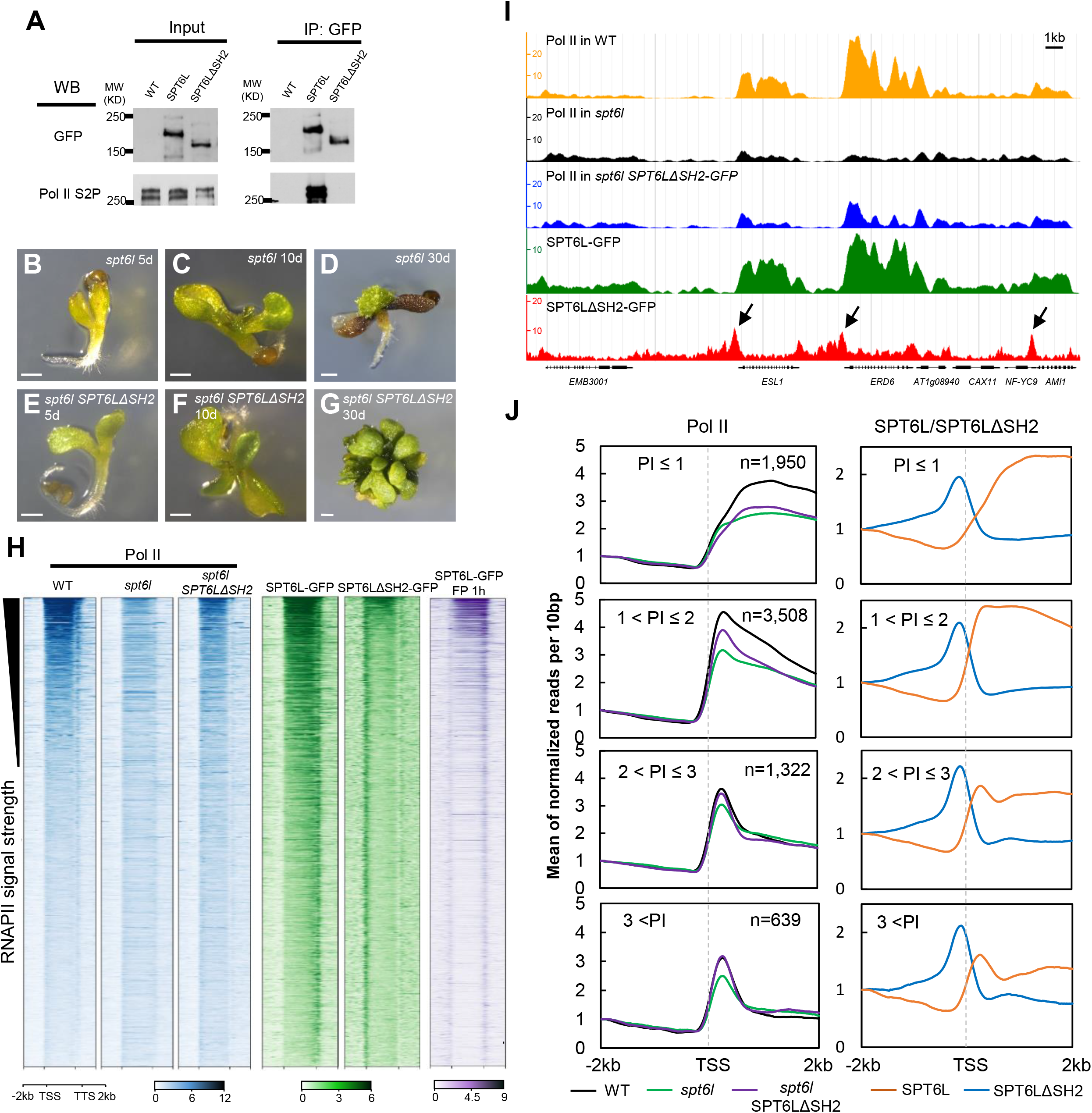
SPT6L enriched at TSS in an RNAPII independent manner. (A) Co-immunoprecipitation performed with transgenic plants expressing SPT6L-GFP and SPT6LΔSH2-GFP, respectively. Antibodies used for IP and immunobloting are indicated. (B) to (D) 5, 10, and 30 days old *spt6l* seedlings, respectively. Bar = 0.5 mm (E) to (G) 5, 10, and 30 days old *spt6l SPT6LΔSH2* seedlings, respectively. Bar = 0.5 mm (H) Heatmaps of RNAPII, SPT6L, and SPT6LΔSH2 binding as measured by ChIP-seq in wide-type (WT), *spt6l*, and *spt6l SPT6LΔSH2* backgrounds, over the same regions shown in Figure 1D. From top to bottom, the plotted genomic regions were sorted by RNAPII signal strength in WT and the values were the means of normalized reads (1x sequencing depth normalization) per 10 bp non-overlapping bin, averaged over two replicates or one replicate for RNAPII in *spt6l*. (I) Screenshots of representative peaks in chromosome 1 visualized in a genome browser (ENPG, www.plantseq.org). The y-axis values represent mean of normalized reads (1x sequencing depth normalization) per 10 bp non-overlapping bin. (J) Mean density of RNAPII, SPT6L, and SPT6LΔSH2 occupancy at the SPT6L bound genes, which were grouped into four groups based on their RNAPII pausing index (PI). Gene number in each group is indicated. The y-axis values were the means of normalized reads (1x sequencing depth normalization) per 10 bp non-overlapping bin, averaged over two replicates or one replicate for RNAPII in *spt6l.* Reads were plotted on regions covering 2kb upstream and downstream of TSS.

We next asked whether the truncated SPT6L can still be recruited to chromatin. To answer that question, we profiled the genome-wide occupancy of SPT6LΔSH2 in *spt6l* mutants. In contrast to the occupancy pattern observed for the full-length SPT6L that spread over the entire transcribed regions of genes, unexpectedly, the signals of SPT6LΔSH2 were found to be enriched at the transcription start sites (TSS) (Figure 2H and 2J). This binding pattern is different from the observed occupancy of SPT6ΔSH2 in yeast, where truncated SPT6 can still weakly spread all-over the transcribed regions (Dronamraju et al., 2018; Mayer et al., 2010; Sdano et al., 2017). Given the morphologic differences between the *spt6l SPT6LΔSH2* and WT seedlings, one can argue that the unexpected TSS enrichment of SPT6LΔSH2 might be caused by altered cell size and/or chromatin structure. To rule out this possibility, we performed a ChIP-seq analysis of SPT6LΔSH2 in *spt6l*^+/-^ and the same pattern was detected again (Figure S2B). As the TSS enrichment of SPT6LΔSH2 was not detected for full-length SPT6L, one can also argue that the enrichment pattern of the truncated SPT6L might result from certain protein structural changes. To assess this possibility, we sought to disrupt the interaction between SPT6L and RNAPII by inhibiting the phosphorylation of RNAPII CTD with flavopiridol (FP), which can block the kinase activity of positive transcription elongation factor b (P-TEFb) (Chao and Price, 2001). We First confirmed the FP inhibition of the phosphorylation of RNAPII CTD and its subsequent effects on plant growth (Figure S2C and S2D). Then, we profiled the genome-wide occupancy of SPT6L after FP treatment, and found dramatically decreased signals over the transcribed regions and concomitant moderate enrichment at TSS (Figure 2H). This pharmacological study complements our genetic work and together they strongly suggest that SPT6L can be targeted to TSS in the absence of its interaction with RNAPII.

The unexpected TSS enrichment of SPT6L drew our attention to its potential effects on RNAPII occupancy around TSS. As the levels of RNAPII around TSS are determined by the equilibrium between its entry and release, we thought that it would be better to take RNAPII pausing at promoter-proximal regions into account. Therefore, we calculated the pausing index (PI) according to a published formula (Gilchrist et al., 2010) and divided RNAPII binding genes into four groups according to their PI values. By plotting RNAPII signals around TSS in WT and *spt6l*, we found that mutation of *SPT6L* led to decreased RNAPII occupancy levels around TSS in all PI groups (Figure 2J), pointing to a role for SPT6L in early transcription stage. Importantly, the introduction of SPT6LΔSH2 can partially or even completely rescue the RNAPII occupancy around TSS in higher PI groups (Figure 2J). In addition, the SPT6LΔSH2 ChIP signals in all the PI groups peaked immediately upstream of TSS, which were followed by RNAPII signals (Figure 2J), suggesting a possible scenario that the presence of SPT6LΔSH2 may help the entry of RNAPII during transcription initiation.

### The HtH and YqgF Domains Are Required for the TSS Association

We next tried to determine which domain(s) of SPT6L is required for its TSS enrichment. *Arabidopsis* SPT6L contains all the five conserved SPT6 domains plus the plant-specific GW/WG domain (Figure 3A). Because the SH2-deleted version of SPT6L can partially rescue the *spt6l* phenotype and show clear enrichment around TSS, we generated five “double-deletion” constructs by deleting each of the five other domains individually on top of the SH2 deletion and introduced them into *spt6l*^+/-^(Figure 3A). All the five versions of truncated SPT6L were localized in the nuclei, as evidenced by the GFP signals in the transgenic root tips (Figure S3A). Further deletion of either the HtH or YqgF domain can compromise the function of SPT6LΔSH2 as shown by the severe phenotype of these transgenic plants (similar to *spt6l*), while the other three double-deletion mutants remain the same as the *SPT6LΔSH2* single mutant (Figure 3B to 3G). This observation suggests that these two domains may be required for the TSS enrichment of SPT6LΔSH2. In addition, we also examined the protein levels of the mutants and saw comparable levels of the truncated SPT6Ls (Figure 3H), indicating that the compromised phenotype was not due to altered protein levels. Finally, we performed ChIP-seq experiments with the *SPT6LΔSH2ΔHtH* and *SPT6LΔSH2ΔYqgF* plants and found that signals around TSS were dramatically reduced in *SPT6LΔSH2ΔHtH* and undetectable in *SPT6LΔSH2ΔYqgF* (Figure 3I). This result suggests that the HtH and YqgF domains are required for the TSS enrichment of SPT6LΔSH2.

**Figure 3.**
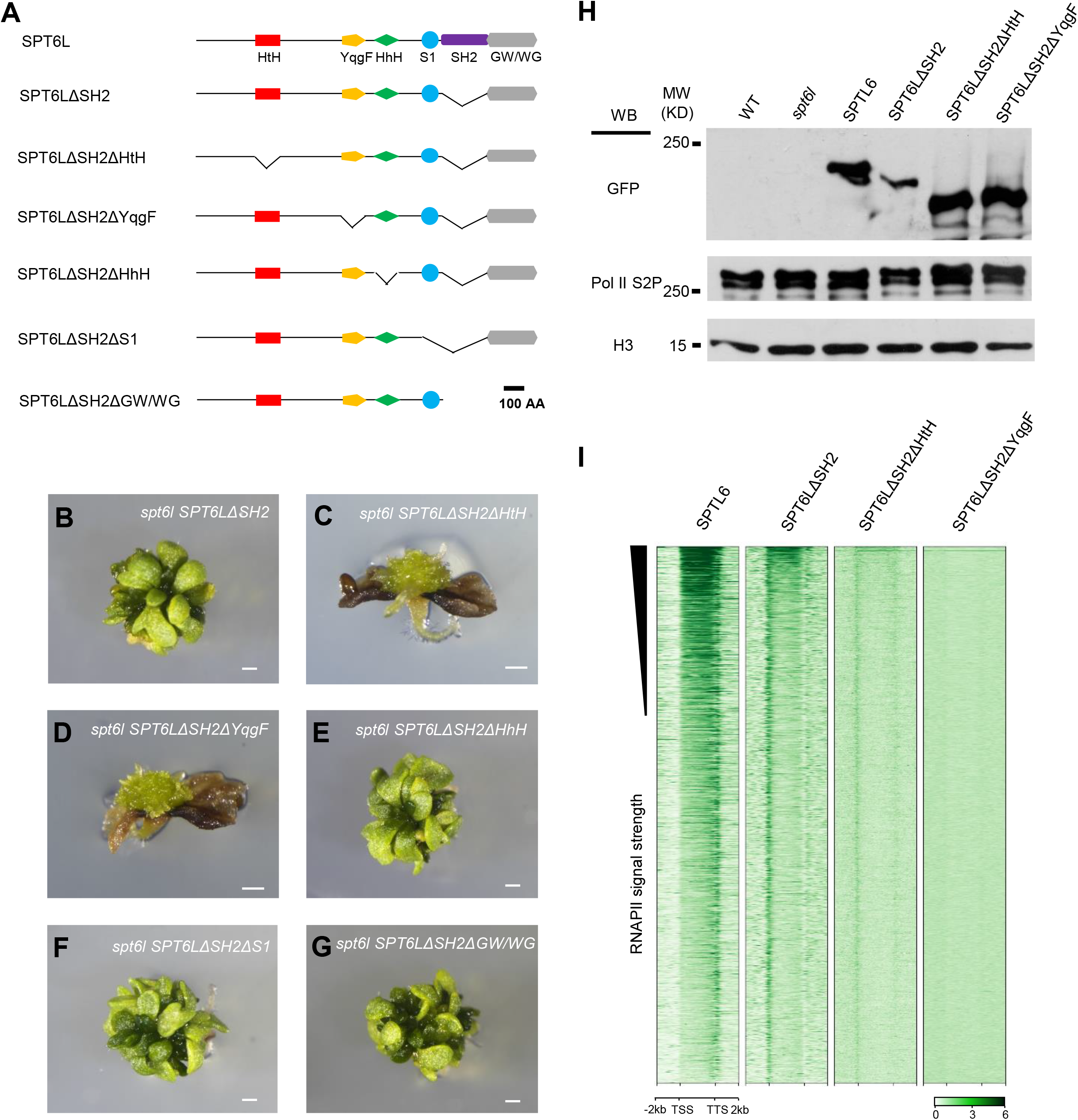
The HtH and YqgF domains are required for the TSS enrichment of SPT6LΔSH2. (A) Diagrams showing the protein domains of Arabidopsis SPT6L and the mutant versions. (B) to (G) 30 days old *spt6l SPT6LΔSH2* (B), *spt6l SPT6LΔSH2ΔHtH* (C), *spt6l SPT6LΔSH2ΔYqgF* (D), *spt6l SPT6LΔSH2ΔHhH* (E), *spt6l SPT6LΔSH2ΔS1* (F), and *spt6l SPT6LΔSH2ΔGW/WG* (G) seedlings, respectively. Bar = 0.5 mm (H) Immunoblots showing the protein levels of truncated versions of SPT6L, and that of RNAPIIS2P in each of the respective genetic backgrounds. H3 levels were used as loading control. (I) Heatmaps of the occupancy of the truncated versions of SPT6L as measured by ChIP-seq, over the same regions and order shown in Figure 2H. The values were the means of normalized reads (1x sequencing depth normalization) per 10 bp non-overlapping bin, averaged over two replicates.

### The HtH and YqgF Domains Are Indispensable for the Distribution of SPT6L Along Genes

We next asked whether the HtH and YqgF domains also contribute to the distribution of SPT6L along transcribed regions. Two new constructs with single deletion of either the HtH or YqgF domain were generated and introduced into *spt6l*^+/-^ plants. The phenotype of the transgenic seedlings indicates that both the single deletion mutants failed to rescue the *spt6l* mutant phenotype (Figure 4A to 4D), which suggests the critical role of HtH and YqgF in maintaining the normal function of SPT6L. To find out how the deletions affect the function of SPT6L in transcription, we first tested whether these two mutant proteins can still interact with RNAPII by performing a Co-IP experiment. As shown in Figure 4SA, they can still interact with RNAPIIS2P, but at a markedly reduced level. We then performed a ChIP-seq analysis to examine their association with chromatin and found that the two versions of truncated SPT6L, although can still weakly associate with RNAPIIS2P, were no longer enriched over gene bodies (Figure 4E). These results indicate that the YqgF and HtH domains are also required for the distribution of SPT6L along genes.

**Figure 4.**
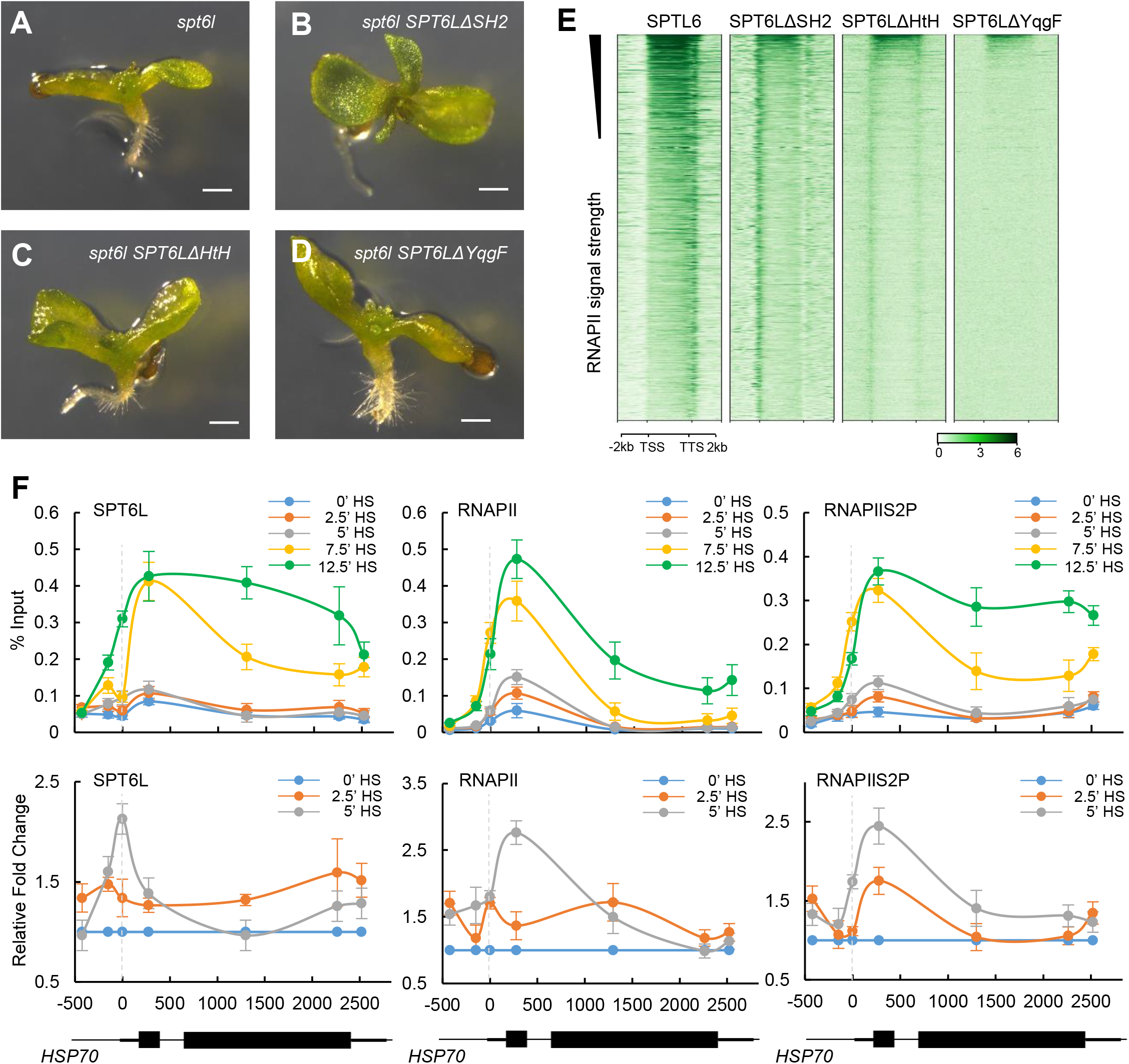
The TSS recruitment of SPT6L precedes its spread along genes. (A) to (D) 10 days old *spt6l* (A), *spt6l SPT6LΔSH2* (B), *spt6l SPT6LΔHtH* (C), and *spt6l SPT6ΔYqgF* (D) seedlings, respectively. Bar = 0.5 mm (E) Heatmaps of the occupancy of the truncated versions of SPT6L as measured by ChIP-seq, over the same regions and order shown in Figure 2H. The values were the means of normalized reads (1x sequencing depth normalization) per 10 bp non-overlapping bin, averaged over two replicates. (F) ChIP signals of SPT6L, RNAPII, and RNAPIIS2P at the *HSP70* locus at different time points after heat shock (HS) treatment. The upper panel shows the ChIP signals relative to the percentage of input. The middle panel shows the fold change of ChIP signals at 2.5 and 5 minutes relative to 0 minutes after HS. Each of the indicated points represents the middle of the PCR fragments. The schematic of the *HSP70* locus is shown in the bottom panel. The plotted values are the means ± S.D. of three biological replicates and numbers on the x-axis are distances to the TSS (TSS = 0).

We next took a genetic approach to examine the functional linkage between the SH2 and HtH/YqgF domains. We reasoned that the *spt6l* phenotype would be rescued in the co-presence of SPT6LΔSH2 and SPT6LΔHtH/SPT6LΔYqgF if the functions of SH2 and HtH/YqgF could be genetically separated. Toward that end, we crossed *spt6l*^+/-^ *SPT6LΔSH2*^+/+^ with either *spt6l*^+/-^ *SPT6LΔHtH*^+/+^ or *spt6l*^+/-^ *SPT6LΔYqgF*^+/+^ and examined the phenotypes of the F1 progenies. Approximately 25% of the F1 progenies (*spt6l*^-/-^ *SPT6LΔSH2*^+/-^ *SPT6LΔHtH*^+/-^ or *spt6l*^-/-^ *SPT6LΔSH2*^+/-^ *SPT6LΔYqgF*^+/-^) showed *SPT6LΔSH2*-like phenotype (Figure S4B to S4D). This result indicates that the functional domains of HtH/YqgF and SH2 have to be preserved in the same SPT6L protein.

### The TSS recruitment of SPT6L precedes its spread over gene bodies

Our genetic and molecular evidence presented above imply that the TSS recruitment of SPT6L may occur prior to its spreading over the gene bodies. To test this hypothesis, we monitored the recruitment of SPT6L and RNAPII at *HEAT SHOCK PROTEIN 70* (*HSP70*, At3g12580) after heat shock (HS) treatment. *HSP70* is a target of SPT6L (Figure S4E) and, as previously reported, its transcription is maintained at a relatively low level at 17°C and dramatically upregulated when temperature elevated to 27°C within 1 hour (Kumar and Wigge, 2010). These features make it a perfect model for investigating the fine dynamics of SPT6L and RNAPII after HS. To optimize the HS condition, we first examined the transcript levels of *HSP70* after HS at 5-minute intervals throughout an hour. Although the level of *HSP70* transcript kept going up throughout the course of the HS treatment, a dramatic change occurred within the first 15 minutes (min) in terms of the increase rate (Figure S4F). Thus, we assessed the occupancy of SPT6L, RNAPII, and RNAPIIS2P at *HSP70* within the first 15 min after HS. As shown in Figure 4F, strong signals of SPT6L were detected downstream of TSS after 7.5 min HS, which is accompanied by the increased signals of RNAPII and RNAPIIS2P at the same sites. After 12.5 min HS, interestingly, we saw further increase in occupancy levels of SPT6L and RNAPIIS2P, but not RNAPII, over the gene body (Figure 4F upper panel), which is consistent with the established role of RNAP II phosphorylation in SPT6L recruitment during elongation. The fact that the level of *HSP70* transcript was increased about 2 fold within the first 5 min after HS (Figure S4F) suggested that the first wave of transcription (after HS) had occurred within that short period of time. Therefore, we re-plotted the ChIP signals of SPT6L, RNAPII, and RNAPIIS2P for the first 5 min after HS. As shown in Figure 4F (middle panel), the SPT6L signals first peaked at TSS after 5 min HS, while both RNAPII and RNAPIIS2P peaked downstream of TSS. This observation suggests that SPT6L was first recruited to TSS and the recruitment was independent of RNAPII in the first wave of transcription at *HSP70* after HS.

## Discussion

Although SPT6 has been extensively studied for its role in transcription elongation, the detailed steps of its recruitment into the transcription machinery have not been elucidated. In this study, we first profiled the genome-wide occupancy of SPT6L and confirmed its conserved function in transcription elongation in plants. Further, we show that SPT6L can bind to transcribing genes at initiation/early elongation regions in an RNAPII independent manner and this binding is indispensable for the loading of SPT6L into the transcription machinery and distribution along gene bodies. Our findings thus have refined the mechanism of SPT6 recruitment and shed light on the roles of SPT6 in transcription initiation.

It has long been observed in yeast that SPT6 plays a role in maintaining the chromatin structure (Ivanovska et al., 2011) and repressing intragenic initiation (Hennig and Fischer, 2013; Kaplan et al., 2003). More recently, during the preparation of this manuscript, a new study further confirmed the role of SPT6 in the repression of intragenic initiation on a genome scale and found reduced genic initiation after knocking out *Spt6* in yeast (Doris et al., 2018). However, it is less clear how Spt6 achieves its roles at initiation sites. Our observation that the TSS enrichment of SPT6LΔSH2 can partially recover the occupancy level of RNAPII in a *spt6l* background (Figure 2H and 2J) point to a role for SPT6 in helping the entry of RNAPII during transcription. Therefore, our result is complementary to the findings in yeast and provides new evidence in support of the role of SPT6 in transcription initiation.

While the recognition between the SH2 domain of SPT6 and the phosphorylated linker region of RNAPII is known to be critical for the recruitment SPT6 over gene body (Sdano et al., 2017), emerging evidence have implied the existence of RNAPII independent recruitment of SPT6 (Adelman et al., 2006; Dronamraju et al., 2018; Mayer et al., 2010; Zhang et al., 2008a). Our work, by integrating genetic and molecular evidence, revealed the RNAPII independent recruitment of SPT6L around TSS region in plants and demonstrate that this recruitment precedes its spreading over the gene body (Figure 4E and 4F). This finding helps to refine the current model of SPT6 recruitment during transcription. Future work on the identification of recruiters of SPT6L at TSS will certainly provide new insight into how SPT6L is involved in transcription initiation and how initiation and elongation are coordinated to ensure a productive transcription.

## Method

### Plant Material and Growth Conditions

The *spt6l* heterozygous seeds (*SALK_016621*) were described previously (Gu et al., 2012) and obtained from the Arabidopsis Biological Resource Center (ABRC) at the Ohio State University.

All Arabidopsis seeds used are in Columbia (Col-0) background. Plants were grown on half strength of Murashige and Skoog (½ MS) medium (0.5XMS salts, 1.5% [w/v] sucrose, and 0.8% agar [pH 5.8]) or soil under 16h/8h light/dark cycle at 23°C. For inhibitor treatment, 5,6-dichloro-1-beta-D-ribofuranosylbenzimidazole (DRB), flavopiridol (FP), or triptolide (Trip) was added at a final concentration of 100, 10, or 10 μM to the media, respectively. For heat shock treatment, seeds were germinated and grown on ½ MS plates for 7 days at 23°C and the plates were then moved to 17°C. After 3 days in 17°C, seedlings were subjected to heat shock treatment at 27°C with different duration. Primers used for genotyping are listed in Supplementary Table 1.

### Plasmid Construction for Plant Transformation

Due to the repetitive nature of the 3’ end sequence of *SPT6L*, we combined PCR amplification and direct DNA synthesis approaches to clone the full-length *SPT6L* genomic region and its 2 kb upstream regulatory sequence. Specifically, part1 (from -2009bp to +6594bp; - and + are relative to ATG) and part2 (from +5247bp to +8443bp) were first PCR-amplified from genomic DNA and cloned into the Gateway entry vector pDONR221 (Invitrogen) by BP reactions. Part3 (from +6889bp to +7380bp, including the repetitive sequence) was synthesized by GenScript (www.genscript.com) and then also inserted into pDONR221. Finally, the entire sequence was assembled by sequential digestions and ligations, first part2-part3 (*AvrII* and *PvuI*) and then part2-3-part1 (*XhoI* and *PvuI*), and cloned into pDONR221 (*ProSPT6L:SPT6L-pDONR221*). Finally, an LR reaction was performed with the destination vector pMDC107(Curtis and Grossniklaus, 2003) to generate the fusion construct with GFP (*ProSPT6L:SPT6L-GFP*). All the domain deletion mutants were generated based on the *ProSPT6L:SPT6L-pDONR221* construct. Primers used are listed in Table S1.

### Analysis of transcript levels

Total RNA extraction, cDNA synthesis, and real-time qPCR were performed as previously described (Chen et al., 2017). Primers used are listed in Table S1. The WT RNA-seq data were obtained from our previous work (Chen et al., 2017). Transcripts were grouped into eight subgroups, from high to low, based on their FPKM values (after conversion to logarithm value (log10)). Finally, the SPT6L ChIP signals were plotted for each of the gene groups separately.

### Co-immunoprecipitation and immunoblot

Seventy-five milligram of 10 day-old seedlings grown on ½ MS medium were homogenized to fine powder in mixer mills and dissolved in 300 μL lysis buffer (20 mM Tris-HCl pH 8.0, 300 mM NaCl, 2.5 mM EDTA, 1 mM DTT, 0.5% TritonX-100, 0.1 mM PMSF, and Protease inhibitor) for 20 minutes at 4°C. Supernatants were collected after centrifuging at x14,000g, 4°C for 10 minutes. For western blot, the supernatants were directly loaded into SDS-PAGE gel. For co-IP, 20 μL anti-GFP μMACS micro-beads (Manufacturer info here) were added into the supernatants and gently shaken at 4°C for 1h. Following the protocol of μMACS GFP isolation kit (130-091-125, MACS), interacting proteins were eluted and loaded into SDS-PAGE gel. Other antibodies used were listed as follows: anti-GFP (ab290, Abcam, lot: GR240324), anti-RNAPII (ab817, Abcam, lot:GR313984), anti-RNAPIISer2P (ab5095, Abcam, lot:GR309257), and anti-Actin (AS13 2640, Agrisera).

### Chromatin immunoprecipitation (ChIP)

ChIP was performed as previously described (Chen et al., 2017) with some modifications. Five grams of 10-day-old *Arabidopsis* seedlings (one gram for *spt6l* and *spt6l SPT6LΔSH2-GFP* seedlings) grown on ½ MS medium were collected. Protein A Dynabeads (Invitrogen) was preincubated with the antibody (5 μL for 50 μL beads) at 4 C in a rotor for at least 6h. After removing excess or unbound antibodies, the pre-cleaned chromatins (cleaned by incubating with Dynabeads alone) were added into antibody bound Dynabeads. To minimize the variations generated from sonication, the same chromatin was equally divided into 2 or 4 tubes and then subjected to different antibodies (2 for anti-GFP and anti-RNAPII in ChIP-seq; 4 for anti-GFP, anti-RNAPII, anti-RNAPIIS2P, and anti-IgG in ChIP-qPCR). ChIP libraries were prepared using the NEBNext^®^ Ultra™ DNA Library Prep Kit (E7370S) following the manufacturer’s instructions and used for Illumina single-end sequencing. Primers used for ChIP-qPCR are listed in Table S1.

### ChIP-seq data analysis

The sequenced reads were aligned to the TAIR10 assembly using the Bowtie2 program (Langmead and Salzberg, 2012) with default settings. After removing unmapped reads and PCR duplicates, peaks were called by using the MACS2 program (Zhang et al., 2008b) with the following setting (-g 135000000, -broad, and -broad-cutoff 0.01). Only the highly reproducible peaks across two biological replicates (IDR ≤ 0.01) were kept. Common genes were identified by using PeakAnalyzer (Salmon-Divon et al., 2010). Coverage files (BigWig files) for all the samples were converted from bam files by using bamCoverage (from deeptools2) (Ramirez et al., 2016) with the following settings (-bs 10 --effectiveGenomeSize 135000000 --normalizeUsing RPGC --ignoreDuplicates -e 300 --samFlagExclude 1796). Heatmaps and mean density plots were generated with deeptools2 (settings indicated in Figure legends). Visualization of coverage files was carried out with a web-based genome browser (ENPG, www.plantseq.org). The genome-wide occupancies of histone methylation and acetylation were obtained from published ChIP-seq datasets (Chen et al., 2017; Luo et al., 2013).

### Data availability

The data that support the findings of this study are available from the corresponding author upon request. The ChIP-seq data have been deposited in Gene Expression Omnibus with the accession code GSE108673.

## ACKNOWLEDGMENTS

We thank the Arabidopsis Biological Resource Centre for providing the mutant seeds used in this study. This work was supported by grants from the Natural Science and Engineering Research Council of Canada (RGPIN/04625-561 2017, to Y.C.); Agriculture and Agri-Food Canada (to Y.C.); Natural Science Foundation of China (31170286, to S.H.); The Specialized Research Fund for the Doctoral Program of Higher Education (20100171110034, to S.H.); National Natural Science Foundation of China (31640059, to J.L.). C.C. and J.S. were supported by a graduate fellowship from the China Scholarship Council.

## AUTHOR CONTRIBUTIONS

C.C. and Y.C. conceived and designed the experiments; J.S. performed immunoblot and Co-immunoprecipitation; R.K.T. performed Real time-qPCR to quantify HSP70 expression level after heat shock; C.L. obtained and validated the *spt6l* mutant. C.C. generated all the constructs, performed ChIP-seq and bioinformatics analyses; V.N., K.Y., Z.Y., S.E.K., J.L., F.M., and S.H. contributed to sequencing, critical reagents, and data analyses. C.C. and Y.C. wrote the paper.

## DECLARATION OF INTERESTS

The authors declare no competing interests.

---

**Figure 1S.**
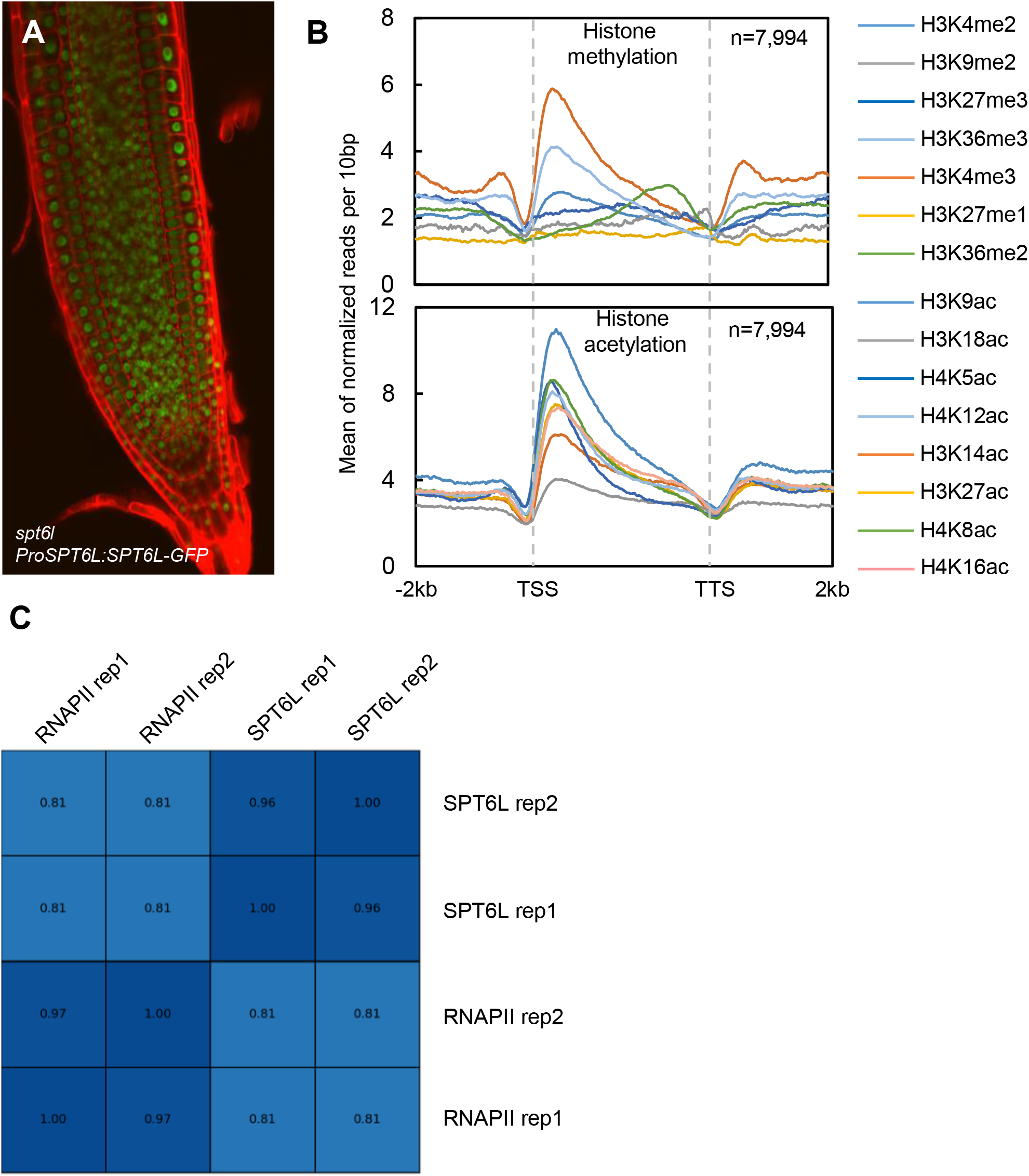
Related to Figure 1 (SPT6L-GFP ChIP-Seq) (A) Root tip (7 DAG) showing nuclear localization of SPT6L. Red signals represent propidium iodide stained cell walls. (B) Mean density of 15 different histone modifications at SPT6L binding genes. Plotting regions are the same as those in Figure 1D, and y-axis values are the means of normalized reads (1x sequencing depth normalization) per 10 bp non-overlapping bin, obtained from one replicate. (C) Reproducibility between ChIP-seq replicates as evaluated by Pearson correlation analysis.

**Figure 2S.**
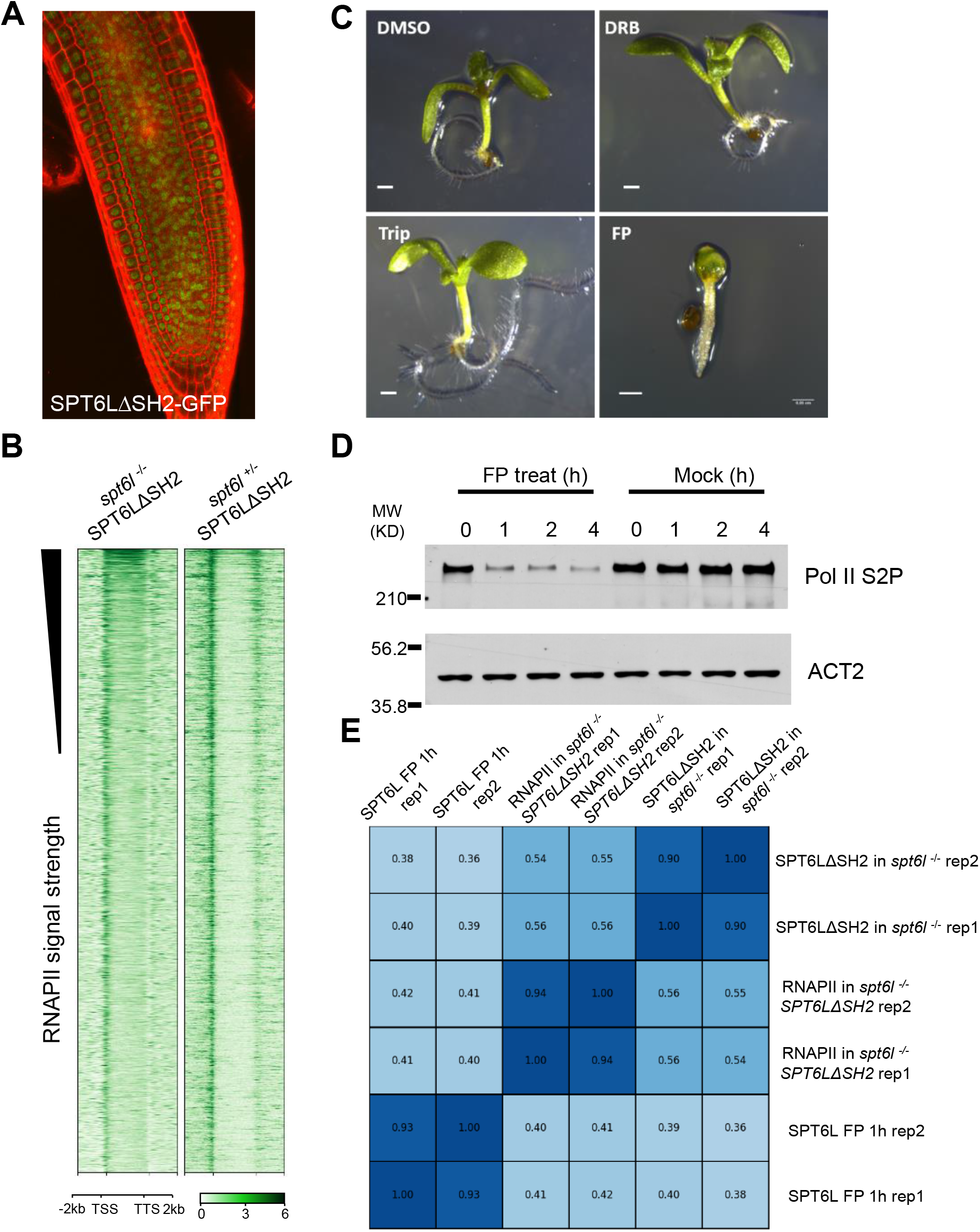
Related to Figure 2 (SPT6LΔSH2 and RNAPIIS2P ChIP-seq) (A) Root tip (7 DAG) showing the nuclear localization of SPT6LΔSH2. Red signals represent propidium iodide stained cell walls. (B) Heatmaps of SPT6LΔSH2 binding as measured by ChIP-seq in *spt6l*^-/-^ and *spt6l*^+/-^ backgrounds, over the same regions and order shown in Figure 2H. The values are the means of normalized reads (1x sequencing depth normalization) per 10 bp non-overlapping bin, averaged over two replicates. (C) Seedlings (10 days old) treated with DMSO, 100 μM 5,6-dichloro-1-beta-D-ribofuranosylbenzimidazole (DRB), 10 μM triptolide (Trip), or 10 μM flavopiridol (FP). Bar = 0.5 mm. (D) Immunoblot showing the level of RNAPIIS2P in 10 days old seedlings after FP treatment for the indicated time. The amount of ACT2 is used as loading control. (E) Reproducibility between ChIP-seq replicates as evaluated by Pearson correlation analysis.

**Figure 3S.**
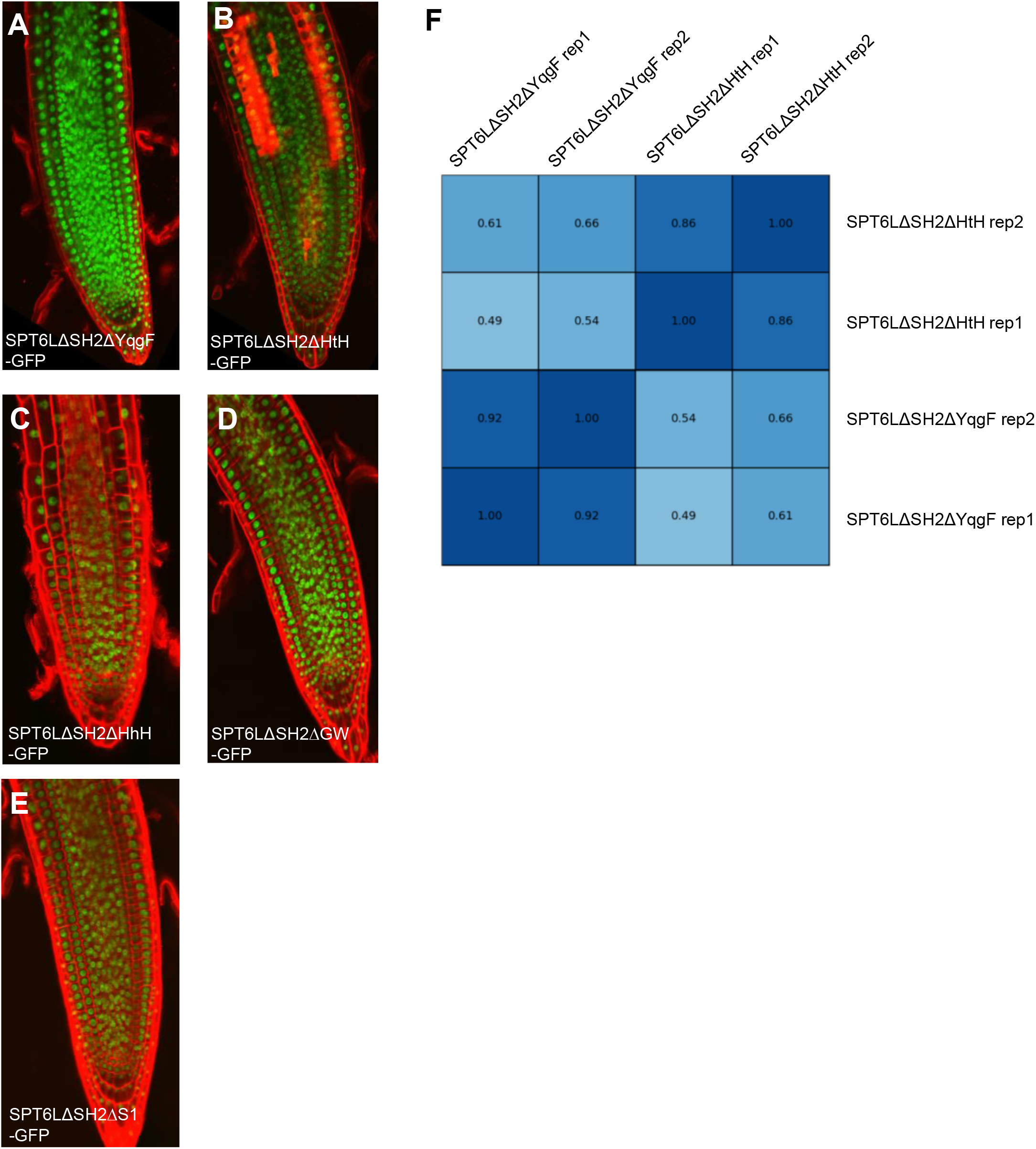
Related to Figure 3 (truncated SPT6Ls and ChIP-seq) (A) to (E) Root tips of 7 days old seedlings showing the nuclear localization of five truncated versions of SPT6L proteins. Red signals represent propidium iodide stained cell walls. (F) Reproducibility between ChIP-seq replicates as evaluated by Pearson correlation analysis.

**Figure 4S.**
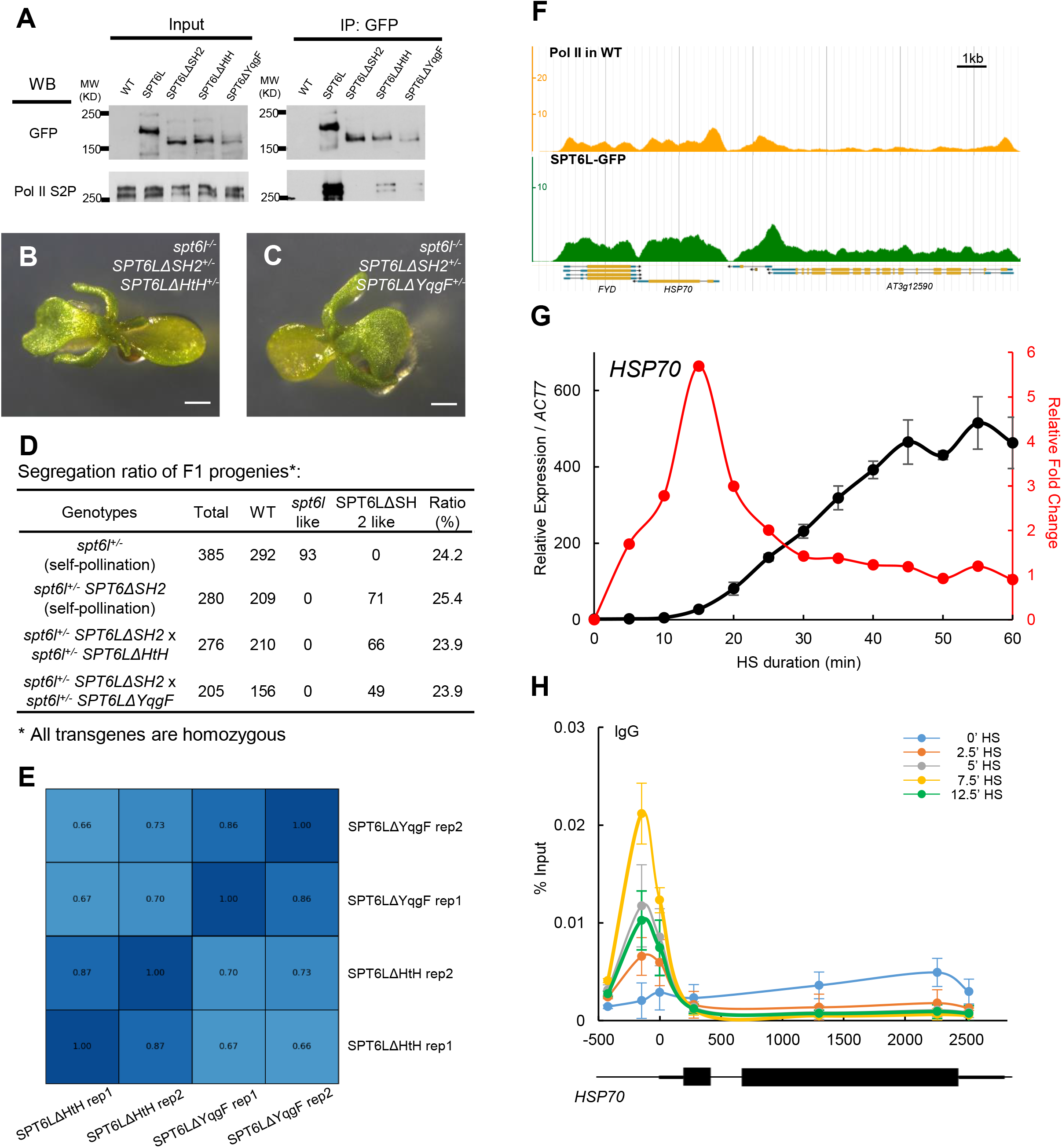
Related to Figure 4 (TSS recruitment of SPT6L precedes its spread along genes) (A) Co-immunoprecipitation examining the interaction between SPT6L-GFP/truncated SPT6Ls and RNAPII. IP and immunoblot performed using indicated antibodies. (B) to (C) F1 progenies (10 DAG) from the cross between *spt6l*^+/-^ *SPT6LΔSH2* with *spt6l*^+/-^ *SPT6LΔHtH* (B) or *spt6l*^+/-^ *SPT6LΔYqgF* (C). Bar = 0.5 mm (D) Segregation ratio of F1 progenies. (E) Reproducibility between ChIP-seq replicates as evaluated by Pearson correlation analysis. (F) Screenshots of RNAPII and SPT6L ChIP-seq signals at the *HSP70* locus in a genome browser (ENPG, www.plantseq.org). The value of y-axis represents mean of normalized reads (1x sequencing depth normalization) per 10 bp non-overlapping bin. (G) The relative transcript levels (left y-axis, black dotted line) and fold change (right y-axis, red dotted line) of *HSP70* during a 1h heat shock (HS) treatment. The relative fold change at each time point was calculated as the ratio between the current and previous time points. The values are the means ± S.D. of three biological replicates. (H) IgG ChIP signals at sites across the *HSP70* gene during a 12.5 minutes heat shock (HS) treatment. Each of the indicated points represents the middle of a PCR fragment; and the schematic of the *HSP70* locus is shown at the bottom. The plotted values are the means ± S.D. of three biological replicates and numbers on the x-axes are distances to the TSS (TSS = 0)

