## Supplementary material for "RNA Polymerase II Independent Recruitment of SPT6 at Transcription Start Sites in *Arabidopsis*": Table S1

Oligos in this work:

| Name | Sequence (5' to 3') | Usage |
| --- | --- | --- |
| LP | TGAAAAACACACAAAATCCAGG | genotyping |
| RP | GATGAGCTTGAGGATGCAAAG | genotyping |
| LB1 | GCGTGGACCGCTTGCTGCAACT | genotyping |
| F1-SPT6L | GGGGACAAGTTTGTACAAAAAAGCAGGCT  TCCTGAAAGACAAACAAGGGTTGATC | SPT6L cloning |
| R1-SPT6L | GGGGACCACTTTGTACAAGAAAGCTGGGT  CCGAGTTACCTTTCCAGCCTCC | SPT6L cloning |
| F2-SPT6L | GGGGACAAGTTTGTACAAAAAAGCAGGCT  TCGGTGAAACTGAGGACACCATA | SPT6L cloning |
| R2-SPT6L | GGGGACCACTTTGTACAAGAAAGCTGGGT  CGAGCTTGATTATGTAATTTAGGTGAA | SPT6L cloning |
| F-ΔHtH | [Phos]CGTTACGAAGAAGAGTCTAG | ΔHtH deletion |
| R-ΔHtH | [Phos]GCTAATTTCATCTACAGGAG | ΔHtH deletion |
| F-ΔYqgF | [Phos]GGTCCAGGTCGGGAAATTTTG | ΔYqgF deletion |
| R-ΔYqgF | [Phos]TTCCTCATCTAAATTAATGTC | ΔYqgF deletion |
| F-ΔHhH | [Phos]CGCAGAAGTGGGCTGGCTGC | ΔHhH deletion |
| R-ΔHhH | [Phos]ACCTGGACCGCATAAAGTTGC | ΔHhH deletion |
| F-ΔS1 | [Phos]GAAAGTGAAATGAGAAACAACAG | ΔS1 deletion |
| R-ΔS1 | [Phos]TATGGTGTCCTCAGTTTCACC | ΔS1 deletion |
| F-ΔSH2 | [Phos]GATGACCCATTGCAAGAATCGGC | ΔSH2 deletion |
| R-ΔSH2 | [Phos]CACATTCTGGTTATGCTGGTGTC | ΔSH2 deletion |
| F-ΔWG/GW | [Phos]GACCCAGCTTTCTTGTACAAAG | ΔWG/GW deletion |
| R-ΔWG/GW | [Phos]AATATGCCTCTGGAAGTATGCCAC | ΔWG/GW deletion |
| qF-ACT7 | CTCATGAAGATTCTCACTGAG | qPCR |
| qF-ACT7 | ACAACAGATAGTTCAATTCCCA | qPCR |
| qF-HSP70 | TCGCCATGAACCCTACCAAC | qPCR |
| qF-HSP70 | CTCACCTGGACCGGAAACAA | qPCR |
| F1-ch-HSP70 | GCAAAAAAAAATCTCAAACACC | ChIP-qPCR |
| R1-ch-HSP70 | CCAAAAAATTTACCCAATTTGAC | ChIP-qPCR |
| F2-ch-HSP70 | AGGACTAAAGAAAAAGTAATTTCC | ChIP-qPCR |
| R2-ch-HSP70 | CAAATAGTAACTCCCGTTTTGAA | ChIP-qPCR |
| F3-ch-HSP70 | CGAAGGAGCTAGAAGCGATAA | ChIP-qPCR |
| R3-ch-HSP70 | TGTTTATATATGAATGAAGAGTGG | ChIP-qPCR |
| F4-ch-HSP70 | AAGGTGAAGGTCCAGCTATCG | ChIP-qPCR |
| R4-ch-HSP70 | CTGTCAGTGAAAGCAACGTAGG | ChIP-qPCR |
| F5-ch-HSP70 | ATTACTGGAACCCGAGAGC | ChIP-qPCR |
| R5-ch-HSP70 | TCCTCGAACCTAGCACGAGT | ChIP-qPCR |
| F6-ch-HSP70 | TGCAATCGACCAAGCTATTG | ChIP-qPCR |
| R6-ch-HSP70 | GTCGTCATCCATTCCTCCTG | ChIP-qPCR |
| F7-ch-HSP70 | TGTCACTCTGAAACTGGTGTGT | ChIP-qPCR |
| R7-ch-HSP70 | TGCCCAGTCGTCTTTCATAGG | ChIP-qPCR |
