## Supplementary material for "RNA Polymerase II Independent Recruitment of SPT6 at Transcription Start Sites in *Arabidopsis*": Table S2

ChIP-seq reads info in this work:

| Sample_name | Raw_reads | Mapped_reads | Unique_reads |
| --- | --- | --- | --- |
| RNAPII_WT_rep1 | 26,067,141 | 24,374,878 | 8,046,844 |
| RNAPII_WT_rep2 | 24,696,082 | 22,968,123 | 8,471,694 |
| WT_Input | 21,611,131 | 21,468,113 | 15,697,258 |
| RNAPII_DSH2_rep1 | 25,129,484 | 22,311,092 | 11,194,276 |
| RNAPII_DSH2_rep2 | 24,795,675 | 19,626,088 | 8,299,877 |
| RNAPII_spt6l | 25,841,440 | 20,509,913 | 9,516,178 |
| spt6l_DSH2_rep1 | 22,531,196 | 15,678,341 | 11,485,810 |
| spt6l_DSH2_rep2 | 21,592,681 | 14,875,174 | 10,902,800 |
| Spt6LD25_Input | 26,137,794 | 25,837,088 | 18,929,234 |
| Spt6LD25_rep1 | 25,332,215 | 25,151,492 | 18,444,455 |
| Spt6LD25_rep2 | 22,666,112 | 22,476,096 | 16,352,341 |
| Spt6LD27_Input | 18,205,420 | 18,054,647 | 13,048,460 |
| Spt6LD27_rep1 | 18,582,373 | 17,732,720 | 14,830,746 |
| Spt6LD27_rep2 | 18,541,191 | 18,035,943 | 14,171,350 |
| Spt6LD2_spt6heterozygous_Input | 13,822,777 | 13,631,610 | 10,030,341 |
| Spt6LD2_spt6heterozygous_rep1 | 18,060,940 | 17,614,712 | 14,285,965 |
| Spt6LD2_spt6heterozygous_rep2 | 18,389,085 | 17,280,883 | 14,396,467 |
| Spt6LD5_Input | 19,940,146 | 19,763,633 | 14,741,134 |
| Spt6LD5_rep1 | 16,935,948 | 16,424,959 | 13,692,287 |
| Spt6LD5_rep2 | 17,717,978 | 17,385,571 | 13,829,683 |
| Spt6LD7_Input | 20,695,317 | 20,511,574 | 14,913,297 |
| Spt6LD7_rep1 | 19,430,121 | 18,724,276 | 16,014,710 |
| Spt6LD7_rep2 | 19,215,065 | 18,785,786 | 15,048,316 |
| Spt6L_DSH2_Input | 25,363,199 | 25,085,269 | 18,321,819 |
| SPT6L_GFP_rep1 | 25,820,848 | 22,842,855 | 18,130,923 |
| SPT6L_GFP_rep2 | 27,393,178 | 25,042,774 | 19,510,516 |
| Spt6LMock_Input | 29,128,493 | 28,874,722 | 20,938,787 |
| Spt6LMock_rep1 | 16,276,584 | 16,019,986 | 13,853,278 |
| Spt6LMock_rep2 | 31,040,844 | 30,562,025 | 24,558,452 |
| Spt6LTreat_Input | 31,322,337 | 31,046,682 | 22,500,793 |
| Spt6LTreat_rep1 | 26,978,032 | 26,508,740 | 22,090,213 |
| Spt6LTreat_rep2 | 27,951,091 | 27,401,246 | 22,907,232 |
| spt6l_input | 26,137,785 | 25,837,088 | 18,929,234 |
